# PIN1 Drives Cellular Plasticity and Immune Modulation in Chronic Pancreatitis

**DOI:** 10.1101/2025.05.13.653850

**Authors:** Vidhi M. Shah, Alexandra Bartlett, Canping Chen, Aubry Matter, Alexander Smith, Hayley Zimny, Xiaoyan Wang, Colin J. Daniel, Jenny Eng, Syber Haverlack, Isaac Youm, Motoyuki Tsuda, Nicholas Calistri, Matthew Reyer, Terry K. Morgan, Gabrielle S. Dewson, Trent Waugh, Dove Keith, Zheng Xia, Koei Chin, Brett C. Sheppard, Jonathan R. Brody, Rosalie C. Sears

**Affiliations:** Brenden-Colson Center for Pancreatic Care, Oregon Health & Science University; Department of Surgery, Oregon Health & Science University; Department of Molecular and Medical Genetics, Oregon Health & Science University; Cancer Early Detection Advanced Research (CEDAR) Center, Oregon Health & Science University; Department of Biomedical Engineering, School of Medicine, Oregon Health & Science University; Department of Pathology and Obstetrics and Gynecology, Center for Developmental Health, Oregon Health & Science University; Knight Cancer Institute, Oregon Health & Science University; Department of Cell, Developmental, and Cancer Biology, Oregon Health & Science University; Computational Biology Program, School of Medicine, Oregon Health & Science University

**Keywords:** PIN1, Chronic Pancreatitis, Cellular Plasticity, Immune Modulation, Fibrosis

## Abstract

**Background and Aims:** Chronic pancreatitis (CP) is characterized by inflammation, fibrosis, and acinar-to-ductal metaplasia (ADM). PIN1, known to drive oncogenic signaling and cellular plasticity in cancer, has an unexplored role in CP. This study investigates PIN1’s expression and function in CP pathogenesis using human tissues and mouse models.

**Methods:** PIN1 expression was assessed in human CP tissue microarrays (TMAs) via immunohistochemistry (IHC) and cyclic immunofluorescence (CyCIF). Acute and chronic pancreatitis were induced in wild-type (WT) and PIN1 knockout (PIN1KO) mice using caerulein. Disease progression was monitored histologically, and immune profiling was conducted using flow cytometry. Pharmacological inhibition was performed using a small molecule PIN1 inhibitor-Sulfopin, and effects were evaluated by histology, qPCR, and cytokine analysis. Single-cell RNA sequencing (scRNA-seq) was performed on pancreatic tissues to perform pathway analysis and intercellular communication.

**Results:** PIN1 expression was elevated in human CP tissues, correlating with disease severity and ADM. In mice, both acute and chronic pancreatitis increased PIN1 expression, but only in our chronic PIN1KO mice displayed reduced pancreatic injury, fibrosis, ADM, and modulated immune infiltration. Pharmacological PIN1 inhibition mimicked the protective effects of genetic knockout, dampening inflammatory pathways. scRNA-seq revealed that PIN1 inhibition altered the intercellular communication networks between epithelial, immune, and stromal cells.

**Conclusion:** PIN1 drives cellular plasticity, immune modulation, and disease progression in CP. Targeting PIN1 may offer a therapeutic strategy to mitigate CP.

**Graphical Abstract:** 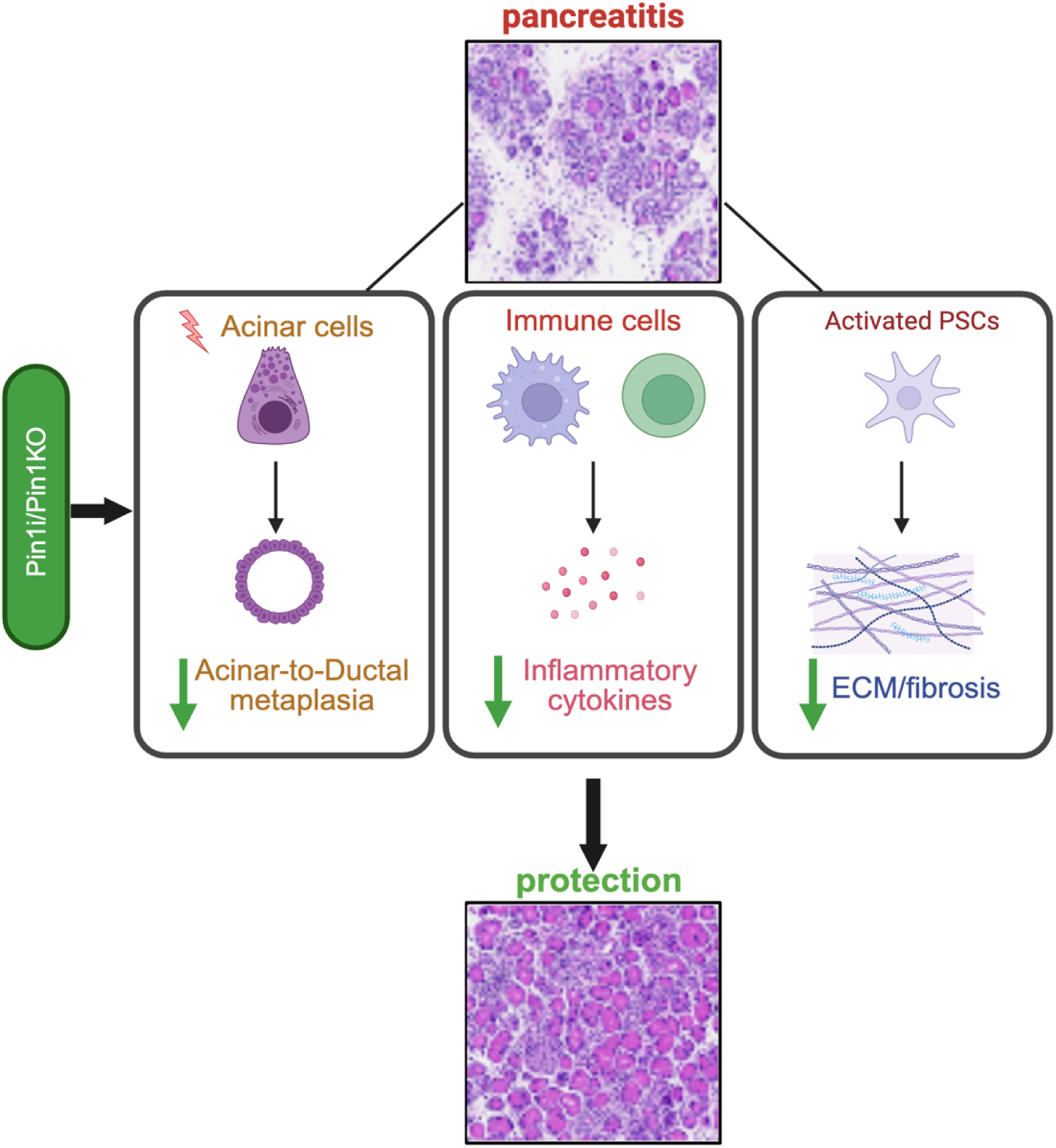

Created in BioRender. Shah, V. (2025) https://BioRender.com/undefined

## Introduction

Chronic pancreatitis (CP) is a debilitating inflammatory disease characterized by progressive acinar cell loss, fibrosis, and acinar-to-ductal metaplasia (ADM), leading to exocrine and endocrine dysfunction and an increased risk of pancreatic ductal adenocarcinoma (PDAC) (PMID: 39647500, PMID: 32798493). While factors such as genetic predisposition, alcohol abuse, and smoking contribute to CP development, the molecular mechanisms underlying its progression remain incompletely understood (PMID: 34461303, PMID: 19308045). A central pathological feature of CP is the plasticity of acinar cells, which undergo ADM-a process involving alterations in gene expression, morphology, and function that increases susceptibility to oncogenic transformation (PMID: 28270694, PMID: 37373094). Chronic inflammation exacerbates this process by recruiting immune and stromal cells, contributing to fibrosis and microenvironment remodeling (PMID: 34915061, PMID: 35922425). Notably, immune cells such as T cells and macrophages drive tissue injury through cytokine production (e.g., TNFα, IL-6) and activation of NF-κB, a master regulator of inflammatory and fibrotic responses (PMID: 37250781, PMID: 19308045). Therefore, identifying key regulators that coordinate epithelial plasticity, immune activation, and stromal remodeling is critical for developing effective therapies.

One such regulator is the peptidyl-prolyl cis/trans isomerase PIN1, which modulates phosphorylation-dependent signaling pathways involved in cell growth, differentiation, and survival (PMID: 36111342, PMID: 38727267). PIN1 is overexpressed in multiple cancers, where it enhances oncogenic signaling by regulating proteins involved in the cell cycle, apoptosis, and epithelial-to-mesenchymal transition (EMT) (PMID: 39418844, PMID: 33537091). Mechanistically, PIN1 isomerizes phosphorylated serine/threonine-proline motifs, altering protein conformation and function (PMID: 33537091). This activity is essential for modulating signaling cascades downstream of growth factors and inflammatory cytokines, including NF-κB and TLR pathways (PMID: 18298383). For instance, PIN1 directly interacts with the p65 subunit of NF-κB, enhancing its nuclear translocation and transcriptional activation of pro-inflammatory genes such as TNFα and IL-18 (PMID: 33537091). In immune cells, PIN1 regulates TLR7/9-dependent type I interferon production by isomerizing IRAK1, enabling IRF7 activation in plasmacytoid dendritic cells (PMID: 21743479). PIN1 also stabilizes AU-rich element-containing mRNAs (e.g., IFN-γ, IL-2) in T cells, amplifying Th1 responses and driving inflammation in autoimmune and allergic disorders (PMID: 17311089, PMID: 17082615). Beyond its roles in inflammation, PIN1 has been implicated in fibrosis across organs. In hepatic stellate cells, PIN1 promotes TGF-β/Smad signaling and collagen production, while in lung fibroblasts, it stabilizes Smad3 to drive extracellular matrix deposition (PMID: 24530597, PMID: 18188456, PMID: 22613712). These findings position PIN1 as a critical integrator of epithelial, immune, and stromal crosstalk-processes central to CP pathogenesis. However, its role in pancreatic injury and remodeling remains unexplored.

Given PIN1’s established roles in epithelial plasticity, NF-κB activation, TLR signaling, and T cell-driven inflammation, and fibrosis -all hallmarks of CP-we hypothesized that PIN1 coordinates multicellular interactions to drive disease progression. To test this, we assessed PIN1 expression in human CP tissues and examined the effects of PIN1 loss-of-function and pharmacological inhibition and performed single-cell RNA sequencing (scRNA-seq) in murine models of pancreatitis. Our study demonstrates that PIN1 is upregulated in CP, where it drives acinar cell injury, ADM formation, immune cell infiltration, and fibrosis. Furthermore, genetic loss or pharmacological inhibition of PIN1 mitigates pancreatic injury, reduces inflammation, immune cell infiltration, and cytokine production, and scRNA-seq shows reduced inflammatory pathways and immune-stromal interactions, highlighting its potential as a therapeutic target in CP. Finally, we integrate scRNA-seq datasets from a published human immune-sorted CP dataset and our caerulein model to establish a PIN1-driven immune signature, revealing conserved pathways that drive disease progression. Together, these findings underscore PIN1 as a key regulator of pancreatic injury and remodeling in CP, providing new insights into the molecular mechanisms that drive chronic inflammation, tissue plasticity, and immune dysfunction.

## Results

### Elevated PIN1 Expression Correlates with Cellular Changes in Human Chronic Pancreatitis

Prior studies have shown that PIN1 is overexpressed in multiple cancers and promotes oncogenic signaling. Additionally, mouse models of breast cancer demonstrate a striking suppression of mammary epithelial duct development in a PIN1-/-background, driven by the regulation of cyclin D1(PMID: 11805292, PMID: 19681904). These findings suggest that PIN1 plays a crucial role in regulating epithelial plasticity. While PIN1’s role in cancer progression has been well-established, its involvement in CP remained unexplored. Our study aimed to address this gap by investigating PIN1 expression in human CP tissues. To address this, we assessed the immunohistochemical (IHC) expression of PIN1 in OHSU-human CP tissue microarrays (TMA) containing 180 cores (1 mm diameter) from 44 unique patient samples: normal background (BG) -CP (4), CP with IPMN (34), PDAC, and 7 control tissues (spleen, placenta, tonsil) (Table 1). Compared to mild chronic pancreatitis cores (Figure 1A), PIN1 expression was significantly increased in severe chronic pancreatitis cores (Figure 1B). Furthermore, PIN1 expression was elevated in ADM lesions within CP cores (Figure 1D) compared to normal BG-CP pancreatic tissue (Figure 1C).To examine PIN1 expression at a single-cell level across different cell types, we employed cyclic immunofluorescence (CyCIF) using a 14-marker panel to assess PIN1 expression in epithelial, endothelial, immune, and stromal cells in both normal and pancreatitis cores. CyCIF staining confirmed significantly higher PIN1 expression in pancreatitis cores compared to normal cores (Figure 1A–B). Moreover, within pancreatitis cores (Figure 1E), PIN1 expression was elevated across all cell types—epithelial, endothelial, immune, and stromal—relative to normal tissue (Figure 1E).

**Figure 1:**
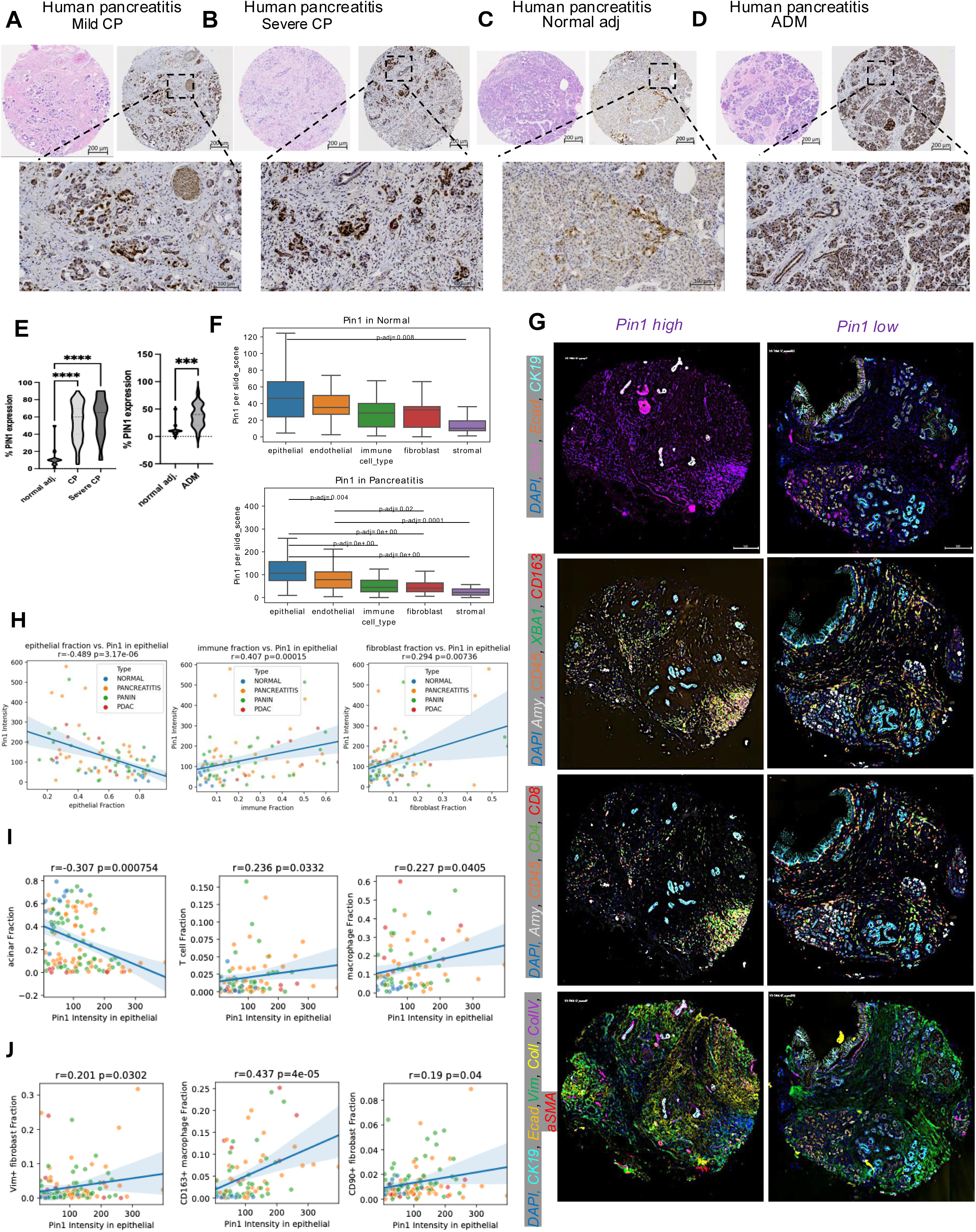
Elevated PIN1 Expression Correlates with Cellular Changes in Human Chronic Pancreatitis A-E. Human pancreatitis specimens were stained and quantified for PIN1 by IHC. **A**. PIN1 expression in the mild/acute pancreatitis (n=29) was compared to **B.** chronic pancreatitis (n=14) samples. **C.** The relative expression of PIN1 in acinar-ductal metaplasia (ADM) (n=11) compared to **D.** adjacent normal (n=5). **F.** Box plot of PIN1 expression in normal (n=4) and pancreatitis (n=32) in different cell types **G**. CycIF staining of representative human pancreatitis microarray (1 mm diameter) core of PIN1 high and PIN1 low of epithelial (amylase, CK19, Ecad) immune (CD45, XBA1, CD163, CD4, CD8), and stromal (Vimentin, Collagens, α-smooth muscle actin) markers, Scale bars = 500 μm. **E**. Correlation between PIN1 intensity in epithelial cells to epithelial, immune, and fibroblast fraction of normal adjacent, pancreatitis, pancreatitis with dysplasia, and PDAC. **F.** Correlation between acinar, T cell, macrophage, vimentin fibroblast, CD163+ macrophage, CD90 + fibroblast – human chronic pancreatitis cellular phenotypes to PIN1 intensity in epithelial cells. t test, mean ± SEM; ∗P ≤ .05, ∗∗P ≤ .01, ∗∗∗P ≤ .001, ∗∗∗∗P ≤ .0001. IHC, immunohistochemistry, CycIF, cyclic immunofluorescence.

**Table 1:** Demographics and Clinical Characteristics.

| Patient Groups | Normal_BG_CP<br>(15 cores)<br>(4 patients) | CP-IPMN<br>(129 cores)<br>(32 patients) | PDAC<br>(12 cores)<br>(6 patients) | Test | P-Value |
| --- | --- | --- | --- | --- | --- |
| % PIN1 expression | 19.7±28.13 | 20.17±21.49 | 20.61±20.62 | Kruskal-Wallis | 0.4354 |
| PIN1 Level:<br>High\|Medium\|Low | 3\|6\|6 | 42\|45\|42 | 5\|5\|2 | Kruskal-Wallis | 0.7330 |
|  | Normal_BG_CP<br>(8 cores)<br>(2 patients) | CP-IPMN<br>(118 cores)<br>(28 patients) | PDAC<br>(13 cores)<br>(6 patients) |  |  |
| Vital Status:<br>Alive/Dead | 8\|0 | 106\|12 | 4\|9 | Kruskal-Wallis | **6.43E-08** |
|  |  | CP-IPMN<br>(52 cores)<br>(13 patients) | PDAC<br>(9 cores)<br>(5 patients) |  |  |
| Survival |  | 2122.1±876.9 | 396.7±202.4 | Mann-Whitney U scaled | **1.96E-06** |
|  | Normal_BG_CP<br>(18 cores)<br>(4 patients) | CP-IPMN<br>(134 cores)<br>(32 patients) | PDAC<br>(13 cores)<br>(6 patients) |  |  |
| Age At Diagnosis | 49.33±8.26 | 68.37±8.11 | 61.38±12.14 | Kruskal-Wallis | **5.28E-09** |
| Sex: Female\|Male | 12\|6 | 74\|60 | 7\|6 | Kruskal-Wallis | 0.6454 |
| Diabetes: No\|DM I\|DM II\|DM - unknown type | 14\|4\|0\|0 | 62\|10\|58\|4 | 9\|10\|4\|4 | Kruskal-Wallis | 0.0675 |
|  | Normal_BG_CP<br>(12 cores)<br>(3 patients) | CP-IPMN<br>(112 cores)<br>(27 patients) | PDAC<br>(13 cores)<br>(6 patients) |  |  |
| HbA1c (%) | 8.3±1.84 | 6.42±1.13 | 6.51±1.66 | Kruskal-Wallis | **0.0009** |

Given the established link between CP and PDAC, along with evidence that pancreatitis increases the pool of KRAS-G12V susceptible acinar precursors (PMID: 17349585) and that pancreatitis-induced tissue damage and inflammation can drive cellular plasticity and transdifferentiation, we further assessed PIN1 expression during disease progression—from normal pancreas to pancreatitis, pancreatic intraepithelial neoplasia (PanINs), and ultimately PDAC. This analysis revealed correlations between PIN1 intensity in epithelial cells and various immune and stromal cell populations (Figure F–H). Specifically, PIN1 expression positively correlated with T cells (CD3+), macrophages (CD68+), M2-like macrophages (CD163+), and fibroblasts (CD90+ and vimentin+) (Figure F–H). In contrast, a negative correlation was observed with acinar cells (amylase+) (Figure F–H), consistent with human disease pathology, where increased immune and fibroblast activity is typically accompanied by a reduction in the acinar (exocrine) compartment of the pancreas. Together, these data suggest that PIN1 expression plays a role in driving the cellular phenotypes of human pancreatitis possibly by exerting a systemic effect through different cell types.

### Elevated PIN1 Expression in Mouse Models of Pancreatitis

To further explore the role of PIN1 in pancreatitis, we assessed its expression in two mouse models: acute and CP (Figure 2A). Wild-type (WT) mice were treated with caerulein—eight hourly injections over two days to induce acute pancreatitis or twice-daily injections for 14 days to model CP (Figure 2A). Compared to normal control pancreas, both caerulein induced-acute and CP resulted in increased PIN1 expression (Figure 2B).

**Figure 2:**
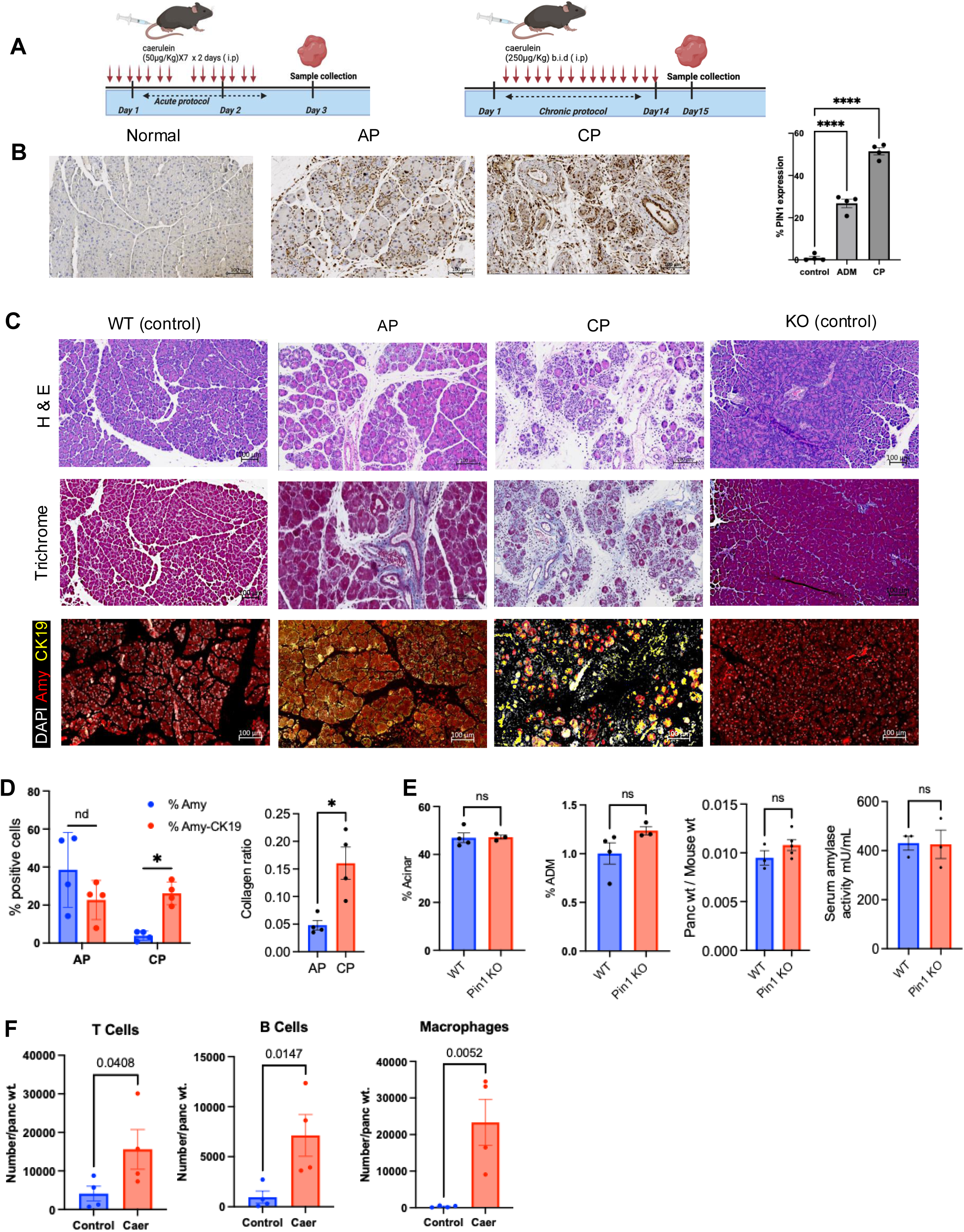
Elevated PIN1 Expression in Mouse Models of Pancreatitis. **A.** Schematic of acute pancreatitis mouse model (mice treated with caerulein (50μg/Kg) hourly for 7 hours daily for 2 days to induce acute pancreatitis (AP) and chronic pancreatitis mouse model (mice treated twice daily with caerulein (250μg/Kg) for 14 days to induce chronic pancreatitis (CP). **B.** Mouse normal tissue, ADM (acute pancreatitis) and CP pancreas tissues were stained and quantified for PIN1 by IHC (n=5). **C**. Representative images for H&E, trichrome, and IF (amylase, CK19, and DAPI) of WT, AP mice, CP mice, and PIN1KO mice **D**. WT and PIN1 KO control mice were quantified for % acinar, %ADM using Visiopharm image analysis software, pancreas weight normalized by animal weight, and serum amylase measurement. **E.** IF quantification of amylase % and ADM (amylase %-CK19%) cells in AP and CP, quantified by quPath. **F.** Flow cytometry of T cells (CD3+), B cells (B220+), macrophages (CD68+) in CP mouse (n=4) compared to control mice (n=4). **G.** Percent collagen deposition measured in control, AP and CP mice (n=5 mice). t test, mean ± SEM; ∗P ≤ .05, ∗∗P ≤ .01, ∗∗∗P ≤ .001, ∗∗∗∗P ≤ .0001. IHC, immunohistochemistry, H&E, hematoxylin and eosin; IF, immunofluorescences; ns, not significant; caer, caerulein.

Histological analysis revealed severe pathological changes with differences in acinar architecture and ductal structures in the chronic models(Figure 2C, 2D). To further characterize these histological changes, we performed co-staining for amylase, amylase-cytokeratin 19 (CK19) co-expressing cells (ADM), and trichrome staining for collagen deposition. The CP model showed a significant increase in ADM-positive cells, a reduction in amylase-expressing cells, and higher collagen deposition compared to the acute caerulein model (Figure 2E).

Interestingly, control whole-body PIN1 knockout (PIN1KO) mice exhibited no significant histological differences in terms of acinar cell percentage, ADM, pancreas weight normalized to body weight, or serum amylase activity compared to WT mice(Figure 2C, 2E). Additionally, immune infiltration, as assessed by flow cytometry, showed increased T cells, B cells, and macrophages in CP compared to control mice (Supplementary Figure 1A). Collectively, these findings demonstrate that the CP caerulein mouse model closely mimics human disease, recapitulating key features such as acinar cell loss, immune infiltration, and increased PIN1 expression.

### PIN1 Deficiency Mitigates Pancreatic Injury and Reshapes Immune Responses in Chronic Pancreatitis

To determine whether PIN1 loss influences acute pancreatic injury, we subjected PIN1KO and WT mice to caerulein treatment following the schematic outlined in Figure 2A. Quantification of acinar and ADM cells from hematoxylin and eosin (H&E)-stained sections using VisioPharm, along with amylase-CK19 co-stained immunofluorescent images analyzed in QuPath, revealed no significant differences between the two groups in the acute caerulein model (Supplementary Figure 2A-C). Additionally, no differences were observed in collagen deposition, pancreas-to-body weight ratios, or the gene expression levels of acinar, ductal, stromal, and inflammatory markers between PIN1 KO and WT mice (Supplementary Figure 2A, 2D-E). Histological analysis at day 7 post-treatment further confirmed that both groups recovered to a normal pancreatic state following the two-day acute protocol (Supplementary Figure 2F).

Given the increased severity of disease presentation in the chronic caerulein model (Figure 2), we next investigated whether PIN1 loss altered disease progression in the two-week protocol (Figure 2A). In contrast to the acute caerulein model, PIN1KO mice exhibited improved retention of acinar cells, as indicated by histological analysis and immunofluorescence staining (Figure 3A-C), reduced collagen deposition, increased body weight, normalized pancreas-to-body weight ratios, and lower serum amylase levels (Figure 3D). Gene expression analysis further demonstrated decreased expression of ductal (Sox9) and stromal (Col1a) markers in caerulein-treated PIN1KO mice compared to WT controls (Figure 3E). To assess immune infiltration, flow cytometry analysis (gating scheme shown in Supplementary Figure 3) revealed a reduction in total T cells and CD8+ cells, along with a trend toward decreased macrophages in PIN1KO mice (Figure 3F, Supplementary Figure 4). While collagen deposition was reduced (Figure 3B), no significant differences were observed in total fibroblast numbers between WT and KO mice following caerulein treatment (Supplementary Figure 4). Additionally, PIN1KO mice displayed increased expression of Tim3 and Tbet, markers associated with a Th1-like immune response, as well as a trend toward higher Foxp3+ CD4+ T cell levels compared to WT mice (Figure 3F).

**Figure 3:**
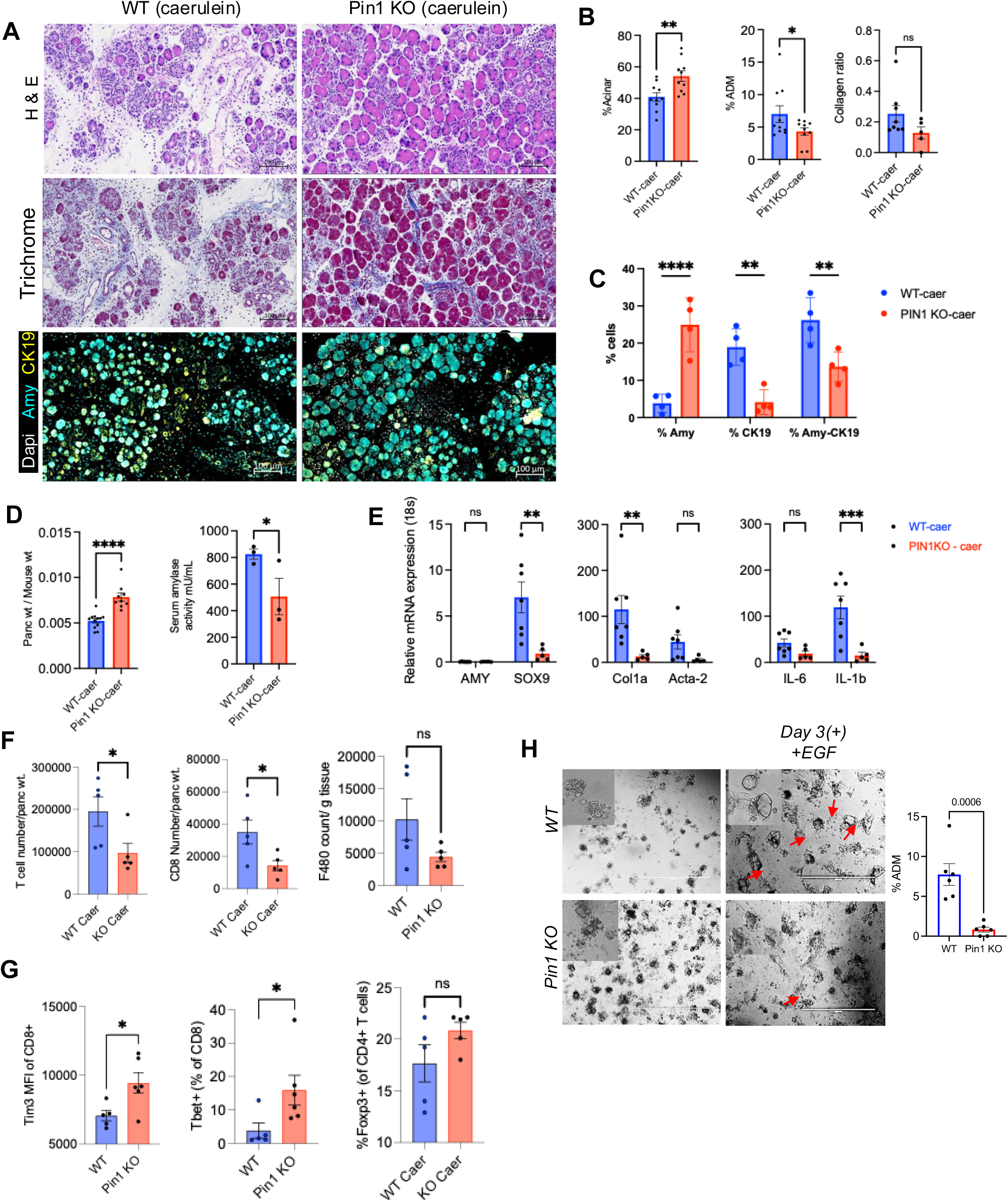
Genetic loss of PIN1 Mitigates Pancreatic Injury and Reshapes Immune Responses in Chronic Pancreatitis. **A.** Representative images for H&E, trichrome, and IF (amylase, CK19, and DAPI) for WT and PIN1KO mice treated with caerulein (250μg/Kg) twice daily for 14 days **B.** % Acinar,% ADM, collagen ratio for control WT and PIN1KO mice**. C.** IF quantification of % amylase, CK19 and amylase-CK19 positive cells. **D.** Pancreas weight normalized by animal weight, serum amylase measurement for control WT and PIN1KO mice. **E.** mRNA expression of amy (acinar), sox9 (ductal), col1a & acta-2 (stromal), Il-6 & Il-1b (pro-inflammatory cytokines) for WT and PIN1KO caerulein treated mice **F.** Flow cytometry expression of T cell count/pancreas weight, CD8+ count/pancreas weight, CD4+ count/pancreas weight. **G.** Tim3 MFI and Tbet (% of CD8), and % Foxp3+ (of CD4+ T cell). **H.** Representative images and quantification of ex-vivo assay of isolated acinar cells from WT and PIN1KO on collagen matrix with EGF. t test, mean ± SEM; ∗P ≤ .05, ∗∗P ≤ .01, ∗∗∗P ≤ .001, ∗∗∗∗P ≤ .0001.; ns, not significant; caer, caerulein, IF, immunofluorescence.

To determine whether these findings persisted in a longer-duration model, we subjected mice to a four-week chronic caerulein protocol (Supplementary Figure 5A). The results mirrored those of the two-week model, with PIN1KO mice exhibiting improved acinar histology, higher pancreas-to-body weight ratios, and reduced collagen deposition compared to WT caerulein-treated mice (Supplementary Figure 5B-C). Similar to the acute caerulein model, both PIN1KO and WT mice showed no differences in recovery at the six-week endpoint (Supplementary Figure 5D).

Additionally, we examined whether PIN1 loss influenced acinar plasticity and ADM formation. Acinar cells isolated from PIN1KO, and WT mice were embedded in a collagen matrix and treated with either PBS or EGF for 72 hours in an ADM assay. Quantification of endpoint cultures demonstrated a significant reduction in ADM structures in PIN1KO acinar cells compared to WT cells (Figure 3G), indicating that PIN1 promotes ADM and acinar cell plasticity in chronic pancreatitis. Together, these findings demonstrate that whole-body loss of PIN1 reduces acinar plasticity, mitigates pancreatic injury, and modulates immune cell subsets in a chronic pancreatitis mouse model, highlighting its systemic role in disease progression.

### Pharmacological Inhibition of PIN1 Mitigates Pancreatic Injury and Modulates Immune Responses in Chronic Pancreatitis

Given the observed increase in PIN1 expression in human and mouse pancreatitis models, along with the reduced injury phenotype in PIN1KO mice, we sought to determine whether pharmacological inhibition of PIN1 using the small-molecule inhibitor Sulfopin (PIN1i) could mitigate chronic pancreatitis progression. To test this, mice were randomized after two weeks of caerulein treatment into two groups: one receiving PIN1i (40 mg/kg) with continued caerulein treatment and the other receiving caerulein alone. Mice were euthanized at the four-week time point, and pancreatic tissues were analyzed (Figure 4A).

**Figure 4:**
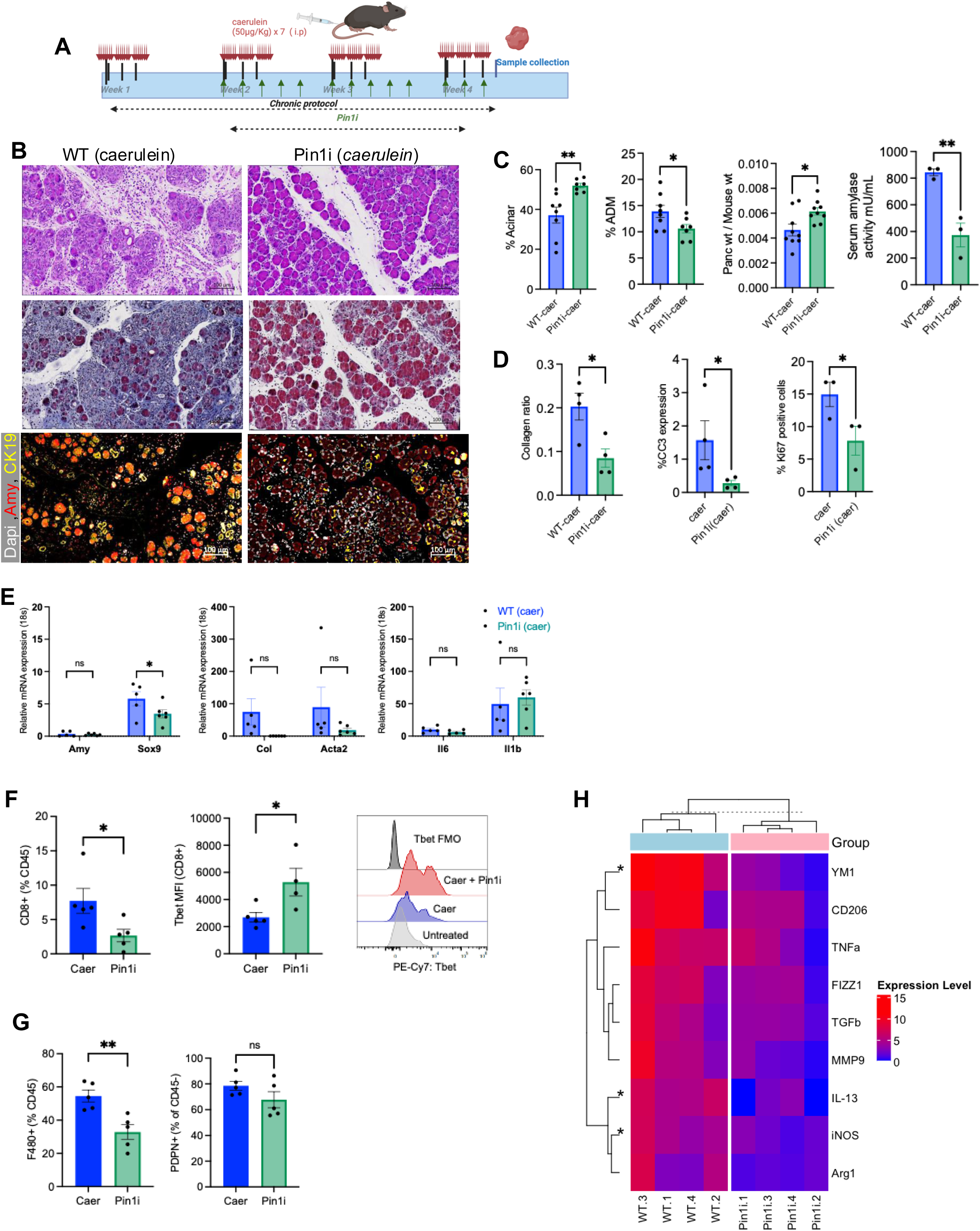
Pharmacological Inhibition of PIN1 Mitigates Pancreatic Injury and Modulates Immune Responses in Chronic Pancreatitis. **A.** Schematics for CP protocol and PIN1i. **B.** Representative images for H&E, trichrome and IF images costained with amylase, CK19, and Dapi to evaluate mouse ADM lesions. **C.** % Acinar (n=8/WT-caer, n=7/PIN1i-caer),% ADM (n=8/WT-caer, n=7/PIN1i-caer), pancreas weight normalized by animal weight (n=9), serum amylase measurement for WT mice treated with caerulein and Pinli (n=3). **D.** Collagen ratio (n=4) and %CC3 expression and %Ki67 in PIN1i+caerulein treated mice compared to caerulein. **E.** mRNA expression of amy (acinar), sox9 (ductal), col1a & acta-2 (stromal), Il-6 & Il-1b (pro-inflammatory cytokines) for WT (n=5 mice) and PIN1i (n=6 mice) caerulein treated mice. **F.** Flow cytometry expression of Tbet MFI of CD8+ and FMO control and %F480 positive cells as % CD45 cells from pancreas of Pinli mice compared to caerulein . **G.** Representative flow cytometry plots and bar graphs depicting macrophages counts per pancreas weight and CD206-MHCII+ (M1) macrophages and CD206+ MFI of F480+ macrophages (M2) from caerulein treated and PIN1i-caerulein treated mice**. H.** mRNA expression of markers of alternatively and classically activated macrophages were assessed using quantitative PCR. Expression of the genes was normalized to their relative expression in control mice for WT and PIN1KO caerulein treated mice (n=4 mice). Data presented as mean ± SEM. t test, 2-way Anova; ∗P ≤ .05, ∗∗P ≤ .01, ∗∗∗P ≤ .001, ∗∗∗∗P ≤ .0001. MFI, mean fluorescence intensity.

PIN1i treatment exhibited no toxicity in normal mice under the same dosing regimen (Supplementary Figure 7). However, in caerulein-treated mice, PIN1i administration resulted in significantly higher acinar cell retention, reduced ADM, an increased pancreas-to-body weight ratio, and lower serum amylase levels compared to untreated caerulein mice (Figure 4B-C). These findings suggest that PIN1 inhibition reduces pancreatic injury, mirroring the protective phenotype observed in PIN1KO mice. Histological analysis, including H&E and trichrome staining, further confirmed an increased acinar cell mass and reduced collagen deposition following PIN1i treatment (Figure 4B). Additionally, PIN1i-treated mice exhibited lower apoptosis (TUNEL assay) and reduced Ki67 expression, indicative of decreased cellular proliferation (Figure 4D).

Gene expression analysis revealed downregulation of ductal (Sox9) and stromal markers (Col1a and Acta2) in PIN1i-treated mice compared to the caerulein-only group (Figure 4E). Immunophenotypic analysis showed a reduction in CD8+ T cells and an increase in Tbet expression, consistent with a shift toward a Th1-like immune response (Figure 4F). Moreover, PIN1i-treated mice exhibited a significant decrease in F4/80+ macrophages as a percentage of CD45+ immune cells compared to caerulein-only mice, although no significant difference was observed in fibroblast populations, as assessed by podoplanin (PDPN) expression (Figure 4G, Supplementary Figure 6).

To further investigate whether PIN1 inhibition affects macrophage polarization, we performed bulk quantitative PCR to assess mRNA expression of markers associated with M1 and M2 macrophage subsets. Compared to caerulein-treated WT mice, the PIN1i-caerulein group displayed reduced expression of both M1- and M2-like markers, with notable decreases in Ym1, IL-13, and iNos expression (Figure 4H, Supplementary Figure 8). Together, these findings demonstrate that pharmacological inhibition of PIN1 reduces acinar cell plasticity, modulates T cell subsets, and alters macrophage polarization in a chronic pancreatitis mouse model. These results further support PIN1 as a key regulator of disease progression and highlight its potential as a therapeutic target for chronic pancreatitis.

### PIN1 Regulates Cytokine and Chemokine Production in Response to Caerulein-Induced Pancreatitis

As we observed no significant differences in T cell activation between the two groups (Supplementary Figure 9A-B) but noted increased expression of both M1 and M2 macrophage markers at the RNA level (Figure 4H), we sought to determine what drives the T cell response and macrophage activity in PIN1-deficient or inhibited mice. To address this, we performed serum cytokine analysis using Luminex (Figure 5). Heatmap analysis (Figure 5A) comparing caerulein-treated PIN1KO and WT mice revealed lower levels of pro-inflammatory cytokines (MIP1b, KC, IFNγ) and T cell-regulating cytokines (IL-2, IL-9, IL-13) in PIN1KO mice. However, other Th1 and Th2 cytokines remained relatively unchanged. Quantitative analysis confirmed that MIP1b and IL-2 levels were significantly reduced in PIN1KO mice compared to WT controls (Figure 5B; p < 0.05). Additionally, volcano plot analysis of serum cytokine gene expression in caerulein-treated mice further validated the significant downregulation of IL-2 and MIP1b in PIN1KO mice relative to WT controls (logFC < −2; p < 0.05). To further examine cytokine production, we performed ex-vivo stimulation of pancreas samples from two-week caerulein-treated mice with IFNγ, IL-2, IL-13, and TNFα. PIN1KO pancreas exhibited a diminished cytokine response compared to WT controls, with a particularly marked reduction in IL-2 levels. This finding suggests impaired T cell differentiation and activation, aligning with the observed serum cytokine profiles. Moreover, we identified CD45+CD3+ cells as the primary source of IL-2 in both WT and PIN1KO mice. Together, these results suggest that PIN1 plays a critical role in driving cytokine production, and its loss leads to a reduction in key pro-inflammatory and T cell-regulating cytokines, potentially altering immune responses in CP.

**Figure 5:**
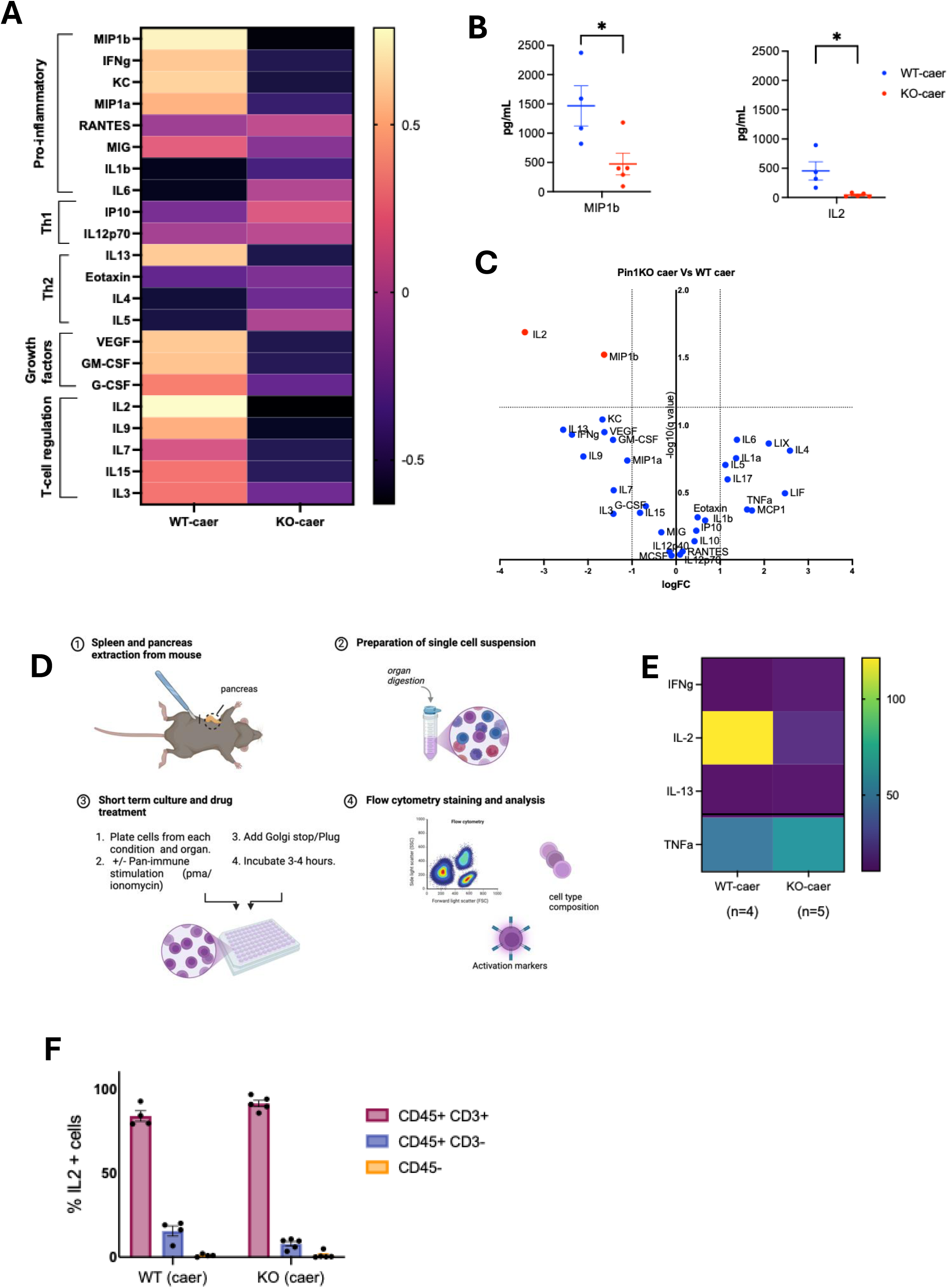
PIN1 Regulates Cytokine and Chemokine Production in Response to Caerulein-Induced Pancreatitis. **A.** Heatmap of cytokines and chemokine expression in serum from caerulein treated PIN1KO (n=5 mice) and WT (n=4 mice) mice quantitated via the Luminex analysis. **B.** MIP1b and IL2 expression in serum of WT-caer and KO-caer mice. **C.** logFC expression of PIN1 KO caer vs WT caer cytokine. **D**. Schematics and quantification for ex-vivo cytokine stimulation assay on pancreas from both WT and PIN1 KO mice treated with caerulein. **E.** % IL-2 secreted from WT-caer and KO-caer mice from different immune cells. Data presented as mean ± SEM. t test, 2-way Anova; ∗P ≤ .05, ∗∗P ≤ .01, ∗∗∗P ≤ .001, ∗∗∗∗P ≤ .0001. caer, caerulein.

### scRNA-seq Reveals PIN1 Inhibition Alters Immune and Cellular Landscapes in Chronic Pancreatitis

Pancreatic tissues from control, caerulein-treated, and PIN1i and caerulein-treated mice were processed for single-cell RNA sequencing (scRNA-seq) using the Evercode cell fixation workflow (Figure 6A). Uniform Manifold Approximation and Projection (UMAP) analysis identified distinct cell populations across conditions (Figure 6B). While overall cell compositions (Figure 6C) were similar among the groups, PIN1i treatment in caerulein-treated mice led to a shift in immune and fibroblast cell distribution compared to caerulein alone (Figure 6C). Differential gene expression analysis demonstrated that caerulein-treated samples exhibited a higher number of upregulated genes in activated macrophages, antigen presenting fibroblast (apFib), and inflammatory fibroblast (iFib) relative to controls, indicating an enhanced inflammatory response (Figure 6D). In contrast, the PIN1i-caerulein group had fewer upregulated genes, suggesting a dampened inflammatory and stromal response compared to caerulein alone. Gene Set Enrichment Analysis (GSEA) using Hallmark pathways further highlighted differences in biological processes between treatment groups (Figure 6E). Compared to control, caerulein treatment significantly enriched pathways associated with “Inflammatory Response,” “TNFa Signaling via NFkB,” “IL6/JAK/STAT3 Signaling,” and “Epithelial-Mesenchymal Transition,” indicating heightened inflammation and epithelial plasticity. However, these pathways were less enriched in the PIN1i-caerulein group, suggesting that PIN1 inhibition attenuates inflammatory signaling. Cell-cell interaction analysis using CellChat revealed distinct intercellular communication networks across groups (Figure 6F). The caerulein-treated group exhibited enhanced interactions between epithelial, immune, and fibroblast populations compared to the control group. Additionally, communication between macrophages, T cells, and fibroblasts was more pronounced in the caerulein group but was significantly reduced in the PIN1i-caerulein condition, indicating that PIN1 inhibition modulates inflammatory cell crosstalk (Figure 6G). Together, these findings suggest that PIN1 inhibition alters the cellular landscape in CP by reducing inflammatory gene expression, modulating immune cell composition, and decreasing cell-cell communication associated with inflammation and fibrosis.

**Figure 6:**
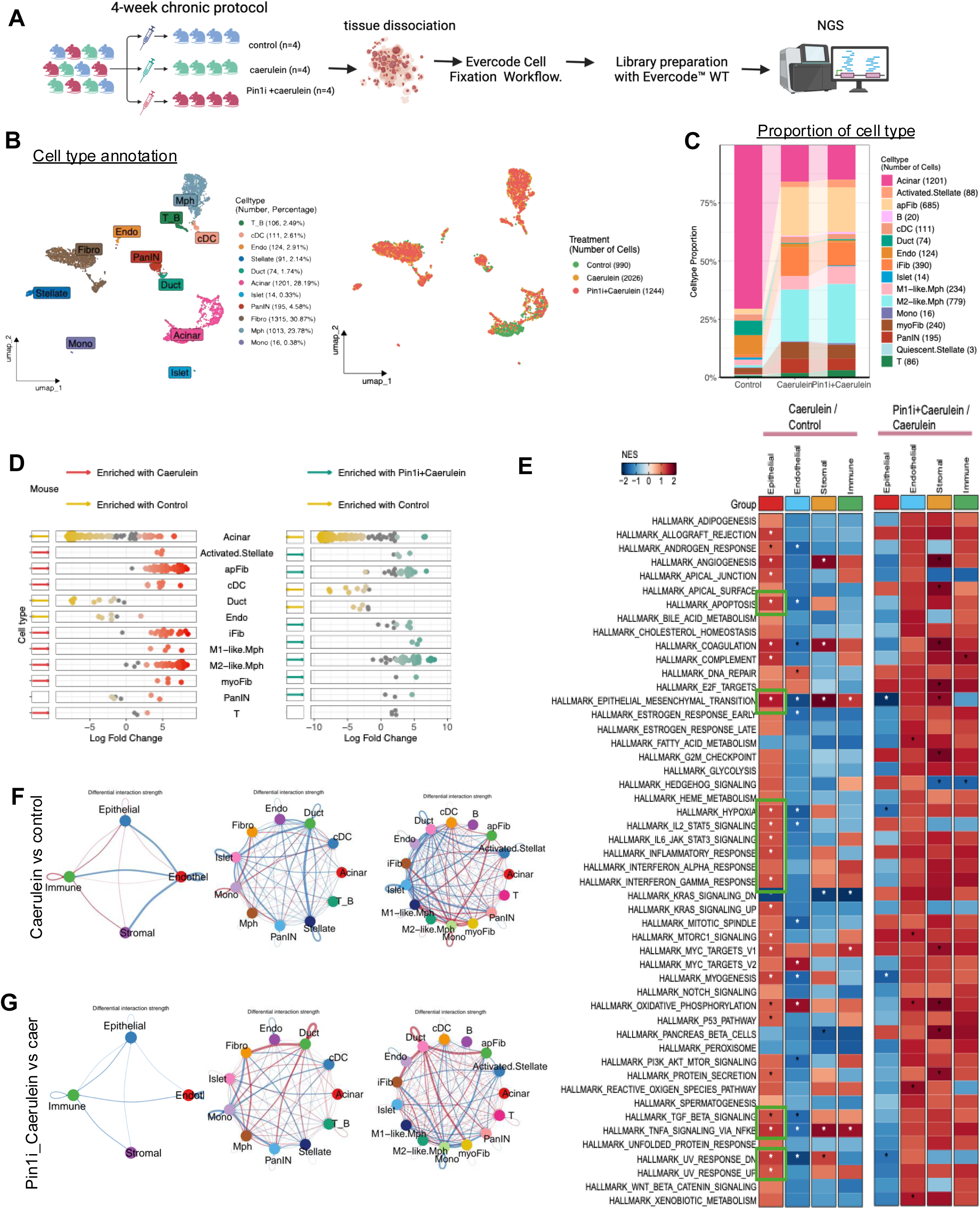
scRNA-seq Reveals PIN1 Inhibition Alters Immune and Cellular Landscapes in Chronic Pancreatitis. **A.** Experimental schematics and workflow of the scRNA-seq (n=4 mice/group). Briefly, mice were treated for caerulein and or PIN1i and collected 12-18h after the last dose of caerulein in week 4 and processed for scRNA-seq analysis using the PARSE workflow **B**. UMAP plots displaying cell populations from all mice colored by major cell type (right plot) and by condition (left plot; control, caerulein, and PIN1i-caerulein) **C.** Bar plot depicting the cellular percentage of major cell type in each of the conditions. **D**. Differentially expressed genes (up-regulated in red; down-regulated in yellow) in caerulein vs control (left) and (up-regulated in green; down-regulated in yellow) PIN1i+caerulein vs control (right). **E.** GSEA Hallmark pathway comparison between caerulein vs control and PIN1i+caerulein vs caerulein groups. **F.** Cell-Chat analysis between caerulein vs control (top) and **G.** PIN1i+caerulein vs caerulein (bottom) groups, showing alterations in cell-cell communication networks.

### Comparative Integration of Human and Mouse Chronic Pancreatitis Reveals PIN1-Driven Immune Signatures

To better understand the transcriptional state of immune cells in human CP and the role of PIN1, we analyzed scRNA-seq data from Lee et al. (PMID: 34702715), which characterizes immune cell dynamics in hereditary and idiopathic CP patients. The analysis of the public human CP dataset revealed distinct PIN1 expression patterns. As shown in Figure 7A, analysis demonstrated significant differences in PIN1 expression across control and experimental conditions (p < 0.0001), with notably higher expression in specific treatment groups (hereditary (Her) and idiopathic (Idio)) compared to controls. While the caerulein-induced mouse model is the gold standard for studying pancreatitis, direct comparisons at the single-cell level between human and mouse models remain limited. Our flow cytometry data (Figure 2) demonstrated increased T cells, B cells, and macrophages in caerulein-treated mice, consistent with observation in human TMA with Cyc-IF (Figure 1). To bridge this gap, we developed a mouse pancreatitis signature by identifying the top 100 differentially expressed genes in caerulein-treated versus control mice. To explore the conservation of immune signatures, we applied the mouse pancreatitis signature to the human dataset. UMAP visualization demonstrated distinct clustering of immune cells, with the mouse pancreatitis signature enriched in human immune cells from PIN1-high samples (Fig. 7B). This cross-species signature application suggests conserved immune mechanisms in pancreatitis pathophysiology. Samples were stratified into PIN1-high and PIN1-low groups based on gene expression levels (Fig. 7C). PIN1-high samples exhibited significantly elevated PIN1 expression compared to PIN1-low samples (p < 0.001), confirming effective stratification. Gene Set Enrichment Analysis (GSEA) comparing PIN1-high versus PIN1-low groups revealed significant enrichment of immune-related pathways in PIN1-high samples (Fig. 7D-E). The PIN1-high group demonstrated robust activation of inflammatory signatures, with prominent upregulation of pathways involved in T cell activation, cytokine signaling, and inflammatory response. Hallmark pathway analysis further revealed enhanced allograft rejection, interferon responses, and complement activation in PIN1-high patients, indicating a pro-inflammatory microenvironment. CellChat analysis revealed increased immune cell-cell communication in PIN1-high patients compared to PIN1-low (Fig. 7F). Key immune subsets, including activated B cells, T helper cells, and macrophages, showed elevated interaction strength, suggesting that PIN1 expression may modulate immune network dynamics during CP. Based on our findings and published role of PIN1 and other drivers of CP, we propose a mechanism (Fig. 7G) by which PIN1 functions as a central amplifier of inflammatory signaling in CP. As illustrated in Fig. 7G, PIN1 could promotes CP through a multifaceted process involving TLR4 signaling (PMID: 38585277), NF-κB activation (PMID: 32347623), and immune cell modulation (PMID: 17082615, PMID: 21743479, PMID: 20956805, PMID: 38851160). Following initial tissue damage, TLR4 activation triggers MyD88-dependent signaling(PMID: 38585277). PIN1 critically enhances this pathway by isomerizing phosphorylated components of the MyD88/IRAK signaling complex, facilitating downstream NF-κB activation(PMID: 32347623). Within this cascade, PIN1 directly interacts with the p65 subunit of NF-κB, promoting its phosphorylation and nuclear translocation(PMID: 32347623). In the nucleus, PIN1 and p65 cooperatively bind to inflammatory gene promoters, enhancing transcription of pro-inflammatory cytokines including TNFα and IL-18(PMID: 32347623, PMID: 38585277). This creates a positive feedback loop where increased cytokine production drives further TLR4 activation and sustained inflammatory signaling. PIN1-mediated amplification of these pathways occurs across multiple cellular compartments - acinar cells experience increased damage and ADM, immune cells (particularly macrophages) shift toward pro-inflammatory phenotypes, and stromal cells adopt activated states promoting fibrosis (PMID: 30660731). To underscores the translational significance of PIN1 as a therapeutic target, we have summarized the findings from our mouse and human data in Table 2. Together, these processes establish a self-sustaining inflammatory environment driving CP progression, positioning PIN1 as a central therapeutic target for interrupting this pathogenic cascade.

**Figure 7:**
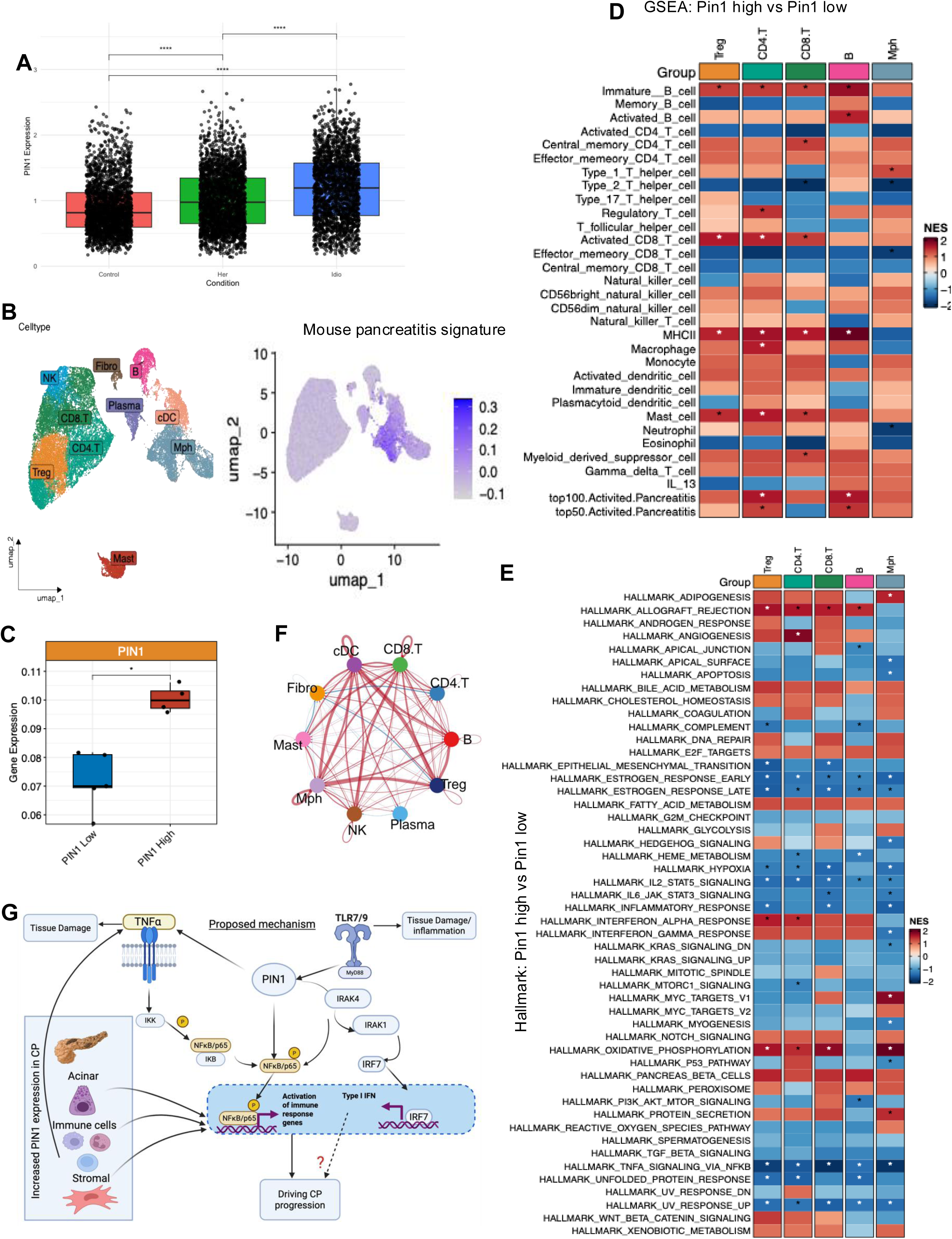
Comparative Integration of Human and Mouse Chronic Pancreatitis Reveals PIN1-Driven Immune Signatures. Single-cell profile integrating mouse and human chronic pancreatitis datasets to investigate PIN1 expression and immune signatures. **A.** PIN1 expression across control and chronic pancreatitis derived from human scRNA-seq data. Patients with CP exhibit significantly higher PIN1 expression compared to controls (p < 0.01). **B.** Mouse pancreatitis signature (top 50 differentially expressed genes from caerulein-treated vs. control mice) applied to human immune cell scRNA-seq data. UMAP visualization of human immune cell subsets annotated by cell type, with overlayed mouse pancreatitis signature scores. Higher pancreatitis scores correspond to increased PIN1 expression, particularly in macrophages, T cells, and fibroblasts. **C.** Chronic pancreatitis patients were stratified into PIN1 high and PIN1 low groups based on PIN1 expression levels (p < 0.05). **D.** Pathway activity comparison between PIN1 high and PIN1 low patient groups across immune cell types in chronic pancreatitis. The heatmap visualization of Gene Set Enrichment Analysis (GSEA) represents NES values for key immune signaling pathways across various immune cell subsets, with red indicating positive enrichment and blue indicating negative enrichment. Asterisks (*) indicate statistically significant enrichment (p < 0.05). The strongest enrichment signals are observed in activated T cell subsets, dendritic cells, and macrophages, suggesting their critical involvement in pancreatitis-associated immune responses. **E.** Cell communication network analysis between PIN1 high and PIN1 low patient groups highlights altered intercellular signaling, including enhanced immune-fibroblast interactions in the PIN1 high group.

**Table 2:** Descriptive elements of the animal model compared to human data.

| Feature | Human Chronic Pancreatitis (Figures) | Mouse Model (Figures) |
| --- | --- | --- |
| PIN1 Expression | Elevated in severe CP (IHC: Figure 1A-B) and ADM lesions (CyCIF: Figure 1D). | Increased in chronic CP models (CyCIF: Fig. 2B). |
| Immune Cell Infiltration | T cells (CD3+), macrophages (CD68+/CD163+), fibroblasts (CD90+/vimentin+) elevated (CyCIF: Figure 1H-G). | Increased T cells, B cells, macrophages in caerulein-treated mice (flow cytometry: Fig. 2F). |
| Disease Severity | Severe CP cores show higher PIN1 expression (IHC: Figure 1A-B). | Chronic caerulein mimics human CP (ADM, fibrosis, immune infiltration)(Fig. 2). PIN1 inhibition reduces severity (Fig.3-4). |
| ADM & Fibrosis | PIN1 elevated in ADM lesions (CyCIF: Figure 1D) and fibrotic regions. | Chronic caerulein induces ADM and collagen deposition. PIN1-KO reduces ADM/fibrosis (histology: Fig. 3). |
| Single-Cell Analysis | scRNA-seq (human immune cells): PIN1-high cells correlate with pancreatitis severity (Figure 7A-C). | scRNA-seq (mouse): PIN1 inhibition alters epithelial-immune-stromal crosstalk (Fig. 6). |
| Immune Pathways | Adaptive cells and MDSC pathways upregulated in PIN1-high immune cells (GSEA: Figure 7D). | TNFα±/NFκB and IL6/JAK/STAT3 pathways (Fig. 6) suppressed in PIN1-KO mice (qPCR/cytokine analysis: Fig.5). |
| Cell-Cell Communication | Enhanced immune-fibroblast interactions in PIN1-high patients (Figure 7F). | PIN1 inhibition reduces pro-inflammatory ligand-receptor networks (CellChat: Fig.6). |
| Adaptive Immune Response | CD4+/CD8+ T and B cells enriched in PIN1-high group (Figure 7D). | PIN1-KO/inhibition reduces effector T cell subsets and shifts macrophage polarization (flow cytometry: Fig. 3-4). |
| Innate Immune Response | Macrophages, dendritic cells, monocytes enriched in PIN1-high group (Figure 7D). | Neutrophils and Tregs reduced in chronic caerulein models (flow cytometry: Fig. 2). |
| Therapeutic Intervention | N/A (observational study). | Pharmacological PIN1 inhibition (Sulfopin) reduces injury/fibrosis (histology/cytokines: Fig. 4-5). |
| Cross-Species Integration | Human immune cells scored against mouse pancreatitis signature (Figure 7B). | Mouse-derived signature validated in human scRNA-seq (Figure 7B). |

## Conclusion

PIN1, a prolyl isomerase, has been widely recognized in the literature as a central regulator of inflammation, fibrosis, and tumor progression across diverse organs. In fibrotic diseases, PIN1 promotes TGF-β/Smad signaling and ECM deposition in the lung and liver. It also modulates immune responses by stabilizing cytokine mRNAs in T cells, facilitating neutrophil chemotaxis, and enabling type I interferon production in dendritic cells. In cancer, particularly PDAC, PIN1 has been shown to enhance tumor growth, metastasis, drug resistance, and immune evasion by acting on tumor cells, cancer-associated fibroblasts, and immune cells.

CP is a complex disease driven by interactions among epithelial, immune, and stromal cells, resulting in persistent inflammation and fibrosis. Motivated by the established roles of PIN1 in these processes in other organs, we investigated its function in CP. Our study demonstrates that PIN1 is highly expressed in human CP tissues, especially in severe cases and areas of acinar-to-ductal metaplasia (ADM), spanning epithelial, stromal, endothelial, and immune compartments. By integrating a mouse model of caerulein-induced pancreatitis with human single-cell RNA sequencing, we identified conserved PIN1-associated mechanisms at single-cell resolution.

Functionally, we show that genetic deletion or pharmacological inhibition of PIN1 in mice reduces pancreatic injury, fibrosis, ADM, and inflammatory cytokine production, and shifts immune responses toward less inflammatory phenotypes.

Taken together, while prior studies have established PIN1 as a key driver of fibrosis, inflammation, and cancer in other organs, our work is the first to demonstrate its critical role in the pathogenesis of CP. Our study highlights PIN1 as a promising therapeutic target in CP and underscores the need for future research to dissect its cell type-specific mechanisms. This will be essential to fully realize the therapeutic potential of PIN1 inhibition in chronic pancreatic disease. Given the central role of T cells, macrophages, and other immune cells in CP pathogenesis, cell-specific knockout models and detailed temporal analyses will be important to delineate PIN1’s direct versus indirect effects on immune cell activation, polarization, and function. Additionally, understanding how PIN1 coordinates epigenetic modifications and metabolic reprogramming in specific immune subsets could reveal novel therapeutic opportunities.

## Methods

### Mouse model

All animal studies were conducted in compliance with Oregon Health & Science University (OHSU) animal use guidelines and were approved by the OHSU Institutional Animal Care and Use Committee (protocol numbers TR1_IP00001014) and *Pin1*^-/-^ mice were maintained on a pure C57/Bl6 background and C57/Bl6 mice were used as WT mice.

### Induction of Pancreatitis

Pancreatitis was induced in age- and litter-matched 8-to 12-week-old mice of both genders. For acute pancreatitis, mice were starved of food overnight, followed by 7 hourly intraperitoneal injections of caerulein (50 μg/kg). Chronic pancreatitis was induced by intraperitoneal injections of caerulein (250 μg/kg) two times daily for 14 days, and the pancreas was collected 16–20 hours later. In additional studies, mice were treated with caerulein three days a week for 4 weeks, every day 7 hourly intraperitoneal injections of caerulein (50 μg/kg) and concurrently Pin1 inhibitor (Sulfopin, Pin1i) (40 mg/Kg) once daily for 3 weeks with caerulein starting 2^nd^ week, pancreas was collected 16-20 hours later.

### Human Pancreatitis Samples

Formalin-fixed paraffin-embedded tissue arrays containing human pancreatitis samples were obtained through the Oregon Pancreas Tissue Registry (IRB00003609) with informed consent. The OHSU IRB approved all protocols, following relevant guidelines. Chronic pancreatitis tissues were collected via fine-needle aspiration (FNA) or Whipple surgery from patients with intraductal papillary mucinous neoplasm (IPMN). Tumor-associated pancreatitis tissues were obtained from PDAC/Intraductal papillary mucinous carcinoma (IPMC) margin resections. The human pancreatitis TMA was generated by a pathologist (T.M) manually marking tissue regions of interest on FFPE sample blocks from surgically resected primary pancreatic tissue samples from patients with pancreatitis, IPMN, or PDAC diagnosed at Oregon Health and Science University. Using a TMA Master II (3DHistech, Hungary) for drilling recipient block and MTA-1 (Estigen Tissue Science, Estonia) for tissue coring, one millimeter tissue cores were punched from representative areas containing normal in background of pancreatitis, chronic pancreatitis in background of IPMN, and PDAC were selected by the pathologist. Clinical and pathological data were retrieved from medical records under institutionally approved protocols. The TMA includes 180 cores from 44 unique subjects (4-chronic pancreatitis, 35-CP in background of IPMN, 6-PDAC, 7-controls (3-spleen. 2-placenta, 2-tonsil)), with each patient contributing 2–6 replicate cores.

### Histology (H&E)/Immunohistochemistry (IHC) and immunofluorescence (IF)

Hematoxylin (Vector Laboratories, Burlingame, CA) and Eosin Y solution (MilliporeSigma) were used for standard H&E staining. Masson’s Trichrome staining was done with Richard-Allan Scientific Masson Trichrome Kit (Thermo Fisher) per manufacturer’s instructions. For immunostains, deparaffinized and rehydrated paraffin-embedded slices. pH9 Tris-EDTA buffer (Dako) (Catlog# pH 9.0) or sodium citrate buffer (Catlog#, pH 6.0) was pressure-boiled for 10 minutes to retrieve antigen. Goat serum and bovine serum albumin were used to quench tissue peroxidase activity and block slices. Primary antibodies were incubated overnight at 4°C on tissue sections: F4/80 (Cell Signaling #70076, 1:800), Ly-6B.2, clone 7/4 (BioRad #MCA771GT, 1:1000), cleaved caspase-3 (Cell Signaling Cat #9664, 1:1000), Ki67 (Abcam #Ab15580, 1:1000). The sections were treated with anti-biotin secondary antibodies (1:500) for 1 hour, the Vectastain ABC kit (Vector Laboratories) for 1 hour, and the DAB substrate for up to 5 minutes for color development. Slides were mounted on Vectamount (Vector Laboratories) after 5 min of hematoxylin counterstaining. For IF, the primary antibodies, amylase (A8273-1VL; 1:100) and CK19 (DSHB, TROMAIII, 1:100) were diluted in blocking buffer and incubated overnight at 4°C. Zeiss AxioScan with Zeiss Zen software captured H&E, Trichrome, and immunostaining pictures (Zeiss Microscopy, Thornwood, NY). QuPath software was used to quantify IF images and on the whole tissue at 20X magnification was analyzed for % acinar and % ADM and collagen ratio with Visiopharm’s software.

### Cyclic immunofluorescence (CycIF)

Immunofluorescence staining, imaging and image processing were performed on the human pancreatitis TMA and mouse pancreatitis tissues as described (PMID: 35545666). Briefly, images were scanned with the Zeiss Axioscan Z1, acquired, stitched and exported to tiff format using Zeiss Zen Blue software (v.2.3), registered using MATLAB (v.9.11.0), followed by cellular segmentation using Cellpose (PMID:33318659) or Mesmer (PMID: 34795433) algorithms. Unsupervised clustering of single cell mean intensity was used to define cell types, using the Leiden algorithm implemented in scanpy (v.1.9.3)(PMID: 29409532). Single-cell mean intensity values of autofluorescence-subtracted images were selected from cytoplasm/nuclear mask for different antibodies based on their cellular distribution as previously published (PMID: 39808504). Briefly, background subtraction was performed by manual thresholding to the total image. Signal was calculated as mean pixel intensity above the threshold, while background was defined as the mean pixel intensity of pixels below the threshold and areas of floating tissue that created bright imaging artifacts and air bubbles that created dark artifacts were manually circled using the napari image viewer and excluded before performing segmentation. The antibodies used for mouse tissue sections are amylase (A8273-1VL; 1:100), CK19 (DSHB, TROMAIII, 1:100), CD45 (ab282747, 1:100), CD4 (50134-R766, 1:100), and CD8 (60168SF, 1:100). The antibodies used on human TMA are Pin1 (G8, 1:100), amylase (A8273-1VL; 1:100), Ki67 (D35B, 1:400), CK19 (A53-B/A2, 1:200), CD8 (C8/468, 1:50), CD4 (EPR6855, 1:100), aSMA (1A4, 1:200), CD68 (KP1, 1:50), CD3 (EPR3094, 1:50), CD45 (EP322Y, 1:50), Vimentin (D21H3, 1:400), and CD163 (EPR14643-36, 1:100). Except amylase, all the antibodies used were conjugated. The Mann-Whitney test and the Kruskal-Wallis test was used to evaluate the correlation between each IF marker. Statistical significance was set at a p-value <0.05.

### Amylase Measurement

Briefly, serum amylase in the blood samples of Pin1KO and C57BL/6 mice from different study groups was estimated using a quantitative colorimetric amylase assay kit (Cat. No. ab102523, Abcam) as per the manufacturer’s protocol (PMID: 37907853, PMID: 36098401). The α-amylase in the serum cleave the substrate ethylidene-pNP-G7 to produce smaller fragments that were eventually modified by α-glucosidase. This caused the release of a chromophore and was measured at optical density (O.D.) = 405 nm using a 96-well multimode plate reader.

### RNA isolation, cDNA preparation, and quantitative PCR

For RNA isolation, harvested pancreatic tissue (previously stored at -80°C.) was homogenized using tissue shearer in 1ml of trizol reagent (Invitrogen) and RNA was isolated using RNAEasy Plus Mini kit (Qiagen) according to manufacturer’s protocol. The harvested RNA was evaluated for quality and concentration using a nanodrop and cDNA was prepared from 2µg of RNA using the Multiscribe Reverse Transcriptase kit (Thermo Fisher). qPCR analysis was performed with Fast SYBR Green reagent (Thermo Fisher) on a StepOne machine (Applied Biosystems). Primers were validated by performing a standard melt curve analysis and are listed in the Supplemental Table 1.

### Cytokine Quantification

Serum cytokines were measured using a Luminex assay. (Eve Technologies, ON, CA) and ex-vivo pancreatic tissue stimulation assays were performed with IFNγ, IL-2, IL-13, and TNFα to quantify cytokine production with and without brefeldin.

### Dissociation of single cells from mouse pancreas

Pancreas samples were minced using scissors and digested in a digestion buffer composed of Collagenase P (2 mg/mL), DNAse I (0.1 mg/mL), and soyaben trypsin inhibitor (0.1mg/mL) at 37°C. To obtain uniform cell suspensions, the dissociated cells were filtered through a 70µm strainer into PBS and subsequently centrifuged at 300xg for 15 mins at 4°C. After removing the supernatant, the pelleted cells were suspended in Red Blood Cell Lysis Solution (130-094-183, Miltenyi Biotec, USA). After being washed once in PBS, the cells were resuspended in Live Dead Aqua (L34966, Life Technologies, USA) or cells were prepared using the Fixation Kit (Parse Biosciences, Seattle, WA, USA) according to the manufacturer’s protocol. The Parse samples were fixed using the Fixation Kit (Parse Biosciences), counted, and stored at -80 °C as per manufacturer’s protocol. We loaded XXX cells per sample for a total of XXX cells from xx samples (4 samples/ 3 groups for this experiment and remaining XX unrelated samples to utilize the maximum capacity of the kit). The samples were multiplexed and processed using the Parse WT kit as per manufacturer’s protocol. All libraries were sequenced as paired-end 150-bp reads using the Illumina NovaSeq 6000 platform.

### Single-Cell RNA Sequencing Workflow for Mouse Pancreatitis Models

(PMID: 39528918) Pancreatic tissues were collected from 12 mice divided into three experimental groups: control (n=4), caerulein-treated (n=4), and Pin1 inhibitor (Pin1i) plus caerulein-treated (n=4). Caerulein was administered to induce chronic pancreatitis, while the Pin1 inhibitor Sulfopin was co-administered in the Pin1i group. Tissues were dissociated into single-cell suspensions using enzymatic digestion with collagenase and DNase, followed by filtration through a 70-μm cell strainer. Cell suspensions were processed using the Evercode cell fixation workflow to preserve RNA integrity. Single-cell RNA sequencing (scRNA-seq) libraries were prepared using the PARSE platform. Barcoded libraries were sequenced on an Illumina NovaSeq 6000 system. For Parse data, we first generated an indexed genome using the mm10 mouse reference downloaded from Ensembl. Next, we used the split-pipe tool from Parse to process files from each sub-library. The results from each sublibrary were then combined into one gene expression matrix. The gene expression matrices were loaded using the Seurat R package (ref). Initial quality control was performed in R using the Seurat package. A Seurat object was created with a minimum threshold of three cells per gene (CreateSeuratObject(min.cells = 3)), and cells with fewer than 200 detected genes were excluded. Cells with >15% mitochondrial gene expression were removed (percent.mt < 15). Cells with a log10 ratio of genes per UMI >0.65 were retained (log10GenesPerUMI > 0.65). Gene expression data were normalized using SCTransform to account for technical variation. Samples were split by experimental group (SplitObject(Pin1i, split.by = ‘sample’)) and processed individually for doublet detection using DoubletFinder, with doublets estimated as 8.4% of the total cell population (0.084*nrow(curr_so@meta.data)). After doublet removal, datasets from all groups were merged into a single Seurat object. To correct for batch effects, Harmony integration was applied to the merged dataset (IntegrateLayers(Pin1i, method=HarmonyIntegration, k.score=20, k.weight=20, orig.reduction=”pca”, new.reduction=”harmony”)). Principal component analysis (PCA) was performed on Harmony-corrected data to identify key dimensions of variation. Cell clustering was performed using neighbor identification: nearest neighbors were identified based on the first 10 principal components (FindNeighbors(dim = 10)) with a clustering resolution of 1 (FindClusters(resolution = 1)). Uniform Manifold Approximation and Projection (UMAP) was used for dimensionality reduction and visualization (RunUMAP(dim = 10)). Differentially expressed genes (DEGs) between experimental groups and clusters were identified using Seurat’s FindMarkers function. Gene Set Enrichment Analysis (GSEA) was performed on DEGs using the Hallmark pathways database to identify biological processes enriched in each condition. Intercellular communication networks across experimental groups were analyzed using CellChat. Ligand-receptor interactions between epithelial, immune, and stromal cell populations were compared to assess differences in signaling pathways between control, caerulein-treated, and Pin1i-caerulein-treated samples. Statistical significance of differences in cell proportions, gene expression levels, and pathway enrichment scores across experimental groups was determined using Wilcoxon rank-sum tests or ANOVA where appropriate. Adjustments for multiple comparisons were made using the Benjamini-Hochberg method.

### Single-cell RNA sequencing analysis of human CP specimen

Single-cell RNA sequencing data from human chronic pancreatitis patients was obtained from Lee et al. (PMID: 34702715), representing both hereditary and idiopathic disease etiologies. The raw data was downloaded from the Gene Expression Omnibus (GEO) database. A mouse pancreatitis signature was generated based on the top 50/100 differentially expressed genes (DEGs) in caerulein-treated vs. control animals from our scRNA-seq dataset. Differentially expressed genes were identified using the Seurat package in R, with a threshold of adjusted p-value < 0.05 and absolute log2 fold change > 0.25. The mouse pancreatitis signature (top 50/100 DEGs) was used to calculate a “pancreatitis score” for each cell in the human chronic pancreatitis scRNA-seq dataset. The AUCell package in R was used for this analysis. Human immune cells were identified and clustered based on canonical cell markers using Seurat. Pin1 expression levels were analyzed within these immune cell subsets. Statistical significance was determined using t-tests or ANOVA with appropriate post-hoc tests.

## Notes

### Competing Interest Statement

The authors have declared no competing interest.

### Funding

- Brenden Colson Center of Pancreatic Care, OHSU

